# Mechanism of germination inhibition of *Clostridioides difficile* spores by an aniline substituted cholate derivative (CaPA)

**DOI:** 10.1101/2023.02.16.528851

**Authors:** Christopher Yip, Ernesto Abel-Santos

## Abstract

*Clostridioides difficile* infection (CDI) is the major identifiable cause of antibiotic-associated diarrhea and has been declared an urgent threat by the CDC. *C. difficile* forms dormant and resistant spores that serve as infectious vehicles for CDI. To cause disease, *C. difficile* spores recognize taurocholate and glycine to trigger the germination process. In contrast to other sporulating bacteria, *C. difficile* spores are postulated to use a protease complex, CspABC, to recognize its germinants. Since spore germination is required for infection, we have developed anti-germination approaches for CDI prophylaxis. Previously, the bile salt analog CaPA (an aniline-substituted cholic acid) was shown to block spore germination and protect rodents from CDI caused by multiple *C. difficile* strains.

In this study, we found that CaPA is an alternative substrate inhibitor of *C. difficile* spore germination. By competing with taurocholate for binding, CaPA delays *C. difficile* spore germination and reduces spore viability, thus diminishing the number of outgrowing vegetative bacteria. We hypothesize that the reduction of toxin-producing bacterial burden explains CaPA’s protective activity against murine CDI. Previous data combined with our results suggests that CaPA binds tightly to *C. difficile* spores in a CspC-dependent manner and irreversibly trap spores in an alternative, time-delayed, and low yield germination pathway. Our results are also consistent with kinetic data suggesting the existence of at least two distinct bile salt binding sites in *C. difficile* spores.

## INTRODUCTION

*Clostridium difficile* infection (CDI) is responsible for approximately 25% of all antibiotic-associated diarrheas [1]. In the U.S., over 500,000 CDI cases occur annually, with a mortality rate of up to 6.5% and costs estimated to be $6.3 billion. CDI has been designated as an urgent threat by the CDC [2]. Furthermore, recurrence of symptoms develops in up to 20% of patients. Therapeutic approach for CDI relapses is currently not completely established [3]. The incidence of CDI is further complicated by the appearance of highly resistant and hypervirulent strains [4, 5].

*C. difficile* forms spores that act as infection vehicles for CDI [6]. Spores do not cause disease but germinate into toxin-producing cells [7-12]. Despite the germination requirement to complete its lifecycle and to initiate disease, all sequenced strains of *C. difficile* lack homologs to the classical Ger receptor proteins found in other *Bacilli* and *Clostridia* species [13-16]. Nevertheless, *C. difficile* spores germinate in response to mixtures of bile salts and amino acids [10, 11].

The absence of conventional Ger receptor genes may suggest *C. difficile* Ger receptors are too divergent from other sporulating bacteria [17]. Kinetic analyses of *C. difficile* spore germination suggested the presence of distinct receptors for taurocholate (a secondary bile salt) and glycine (the smallest amino acid) [12].

The *cspABC* operon has been associated with taurocholate-mediated germination of *C. difficile* spores. Specifically, the CspC protease has been implicated as the bile salt germination receptor [8]. However, high abundance of CspC has also been shown to downregulate *C. difficile* spore germination by inhibiting cortex hydrolysis [18]. More recently, the pseudoprotease CspA has been suggested as a second taurocholate receptor [19].

The role of CspC as a *bona fide* germination receptor has been disputed since CspC seems to be involved not only in the detection of bile salts, but also of DPA and amino acids [20]. These new studies suggest that CspC functions as a signaling node rather than a ligand-binding receptor [21].

Strain-to-strain germination variability has been reported in *C. difficile* [22]. Indeed, chenodeoxycholate, a naturally occurring bile salt, moderately inhibits the germination of *C. difficile* type strain 630 [9]. However, when chenodeoxycholate was assayed against a panel of other *C. difficile* clinical isolates, a large range of inhibition efficacies was observed [22].

Furthermore, different strains seem to also have varying responses for the germinant taurocholate [22]. These inter-strain differences in recognizing bile salts have been attributed to CspC working to first activate spore germination and then inhibiting downstream processes [18].

Because of the critical nature of spore germination in the onset of infection, we expect that anti-germination therapy will prevent CDI and its relapse [23]. Given the role that bile salts play in *C. difficile* spore germination, we and others have examined sterane compounds as inhibitors of spore germination [11, 24-32].

We have developed cholates containing *m*-sulfanilic acid (CamSA), or aniline (CaPA) sidechains that inhibit *C. difficile* spore germination at micromolar concentrations even when the taurocholate germinant is present at saturating millimolar levels [11, 33-36]. As our first CDI prophylactics, CamSA and CaPA have been extensively characterized *in vitro* and *in vivo* [27, 33-38].

Consistent with previous bile salt susceptibility data, we recently found that CamSA inhibits *C. difficile* strain 630 spore germination *in vitro* [11], but was less active against spores from other *C. difficile* ribotypes, including the hypervirulent strain R20291 [33]. Strain-specific *in vitro* germination activity of CamSA correlated with its ability to prevent CDI in mice.

In contrast, CaPA was found to be a better anti-germinant than CamSA against eight different *C. difficile* strains [33]. In addition, CaPA reduced, delayed, and/or prevented murine CDI signs from all strains tested. These data suggest that *C. difficile* spores respond to germination inhibitors in a strain-dependent manner.

In this study, we found that anti-germinant selectivity by *C. difficile* strain is not correlated to different CspC variants. We also tested the mechanism of CaPA-mediated inhibition of *C. difficile* spore germination. We found that CaPA is an alternative substrate inhibitor of *C. difficile* spore germination, and thus creates a kinetic bottleneck for spore outgrow into vegetative cells. We further found that CaPA can outcompete taurocholate for spore binding and irreversibly trap spores in an alternative, slowed-down, and low yield germination pathway. In line with previous kinetic data, these results are consistent with the existence of at least two distinct bile salt binding sites in *C. difficile* spores.

## METHODS

### Materials

Taurocholate and glycine were purchased from Sigma-Aldrich Corporation (St. Louis, MO). Bile salt analogs CaPA and CamSA were synthesized and kindly provided by Prof. Steven Firestine at Wayne State University. All bile acid analogs were dissolved in DMSO. Glycine was dissolved in deionized water (DIH2O). Primers were ordered from Integrated DNA Technologies (IDT; Coralville, IA). Genomic DNA was extracted using Wizard Genomic DNA Purification Kit (Promega; Madison, WI). PCR products and amplicons were purified using QIAquick PCR Purification kit (Qiagen; Hilden, Germany). PCRs were conducted using Phusion Hot Start II DNA Polymerase (Thermo Fisher; Waltham, MA). BenchTop 1 kb and 100 bp DNA ladders were purchased from Promega. Other reagents and media were obtained from VWR International (Radnor, PA).

### Bacterial Strains

*C. difficile* strains R20291, 7004578, CDC38, DH1834, 05-1223-046, 8085054, and 9001966 were kind gifts from Prof. Nigel Minton at the University of Nottingham, UK. *C. difficile* strain 630 was obtained from the American Type Culture Collection (ATCC).

Cloning was performed in *E. coli* strain Mach1 2T1 (Invitrogen, Waltham, MA). *E. coli* strain CA434 (*E. coli* HB101 harboring pRK24 and used for conjugations), *C. difficile* strain 630*Δerm ΔpyrE*, and plasmid pMTL-YN3 were kind gifts from Prof. Aimee Shen at Tufts University.

### Spore Preparation and Purification

Frozen stocks of *C. difficile* strains were individually streaked onto pre-reduced BHIS plates (BHI agar supplemented with 0.5% yeast extract, and 0.1% L-cysteine) and incubated overnight at 37 °C in an anaerobic environment (10% CO_2_, 10% H_2_, and 80% N_2_) to yield single colonies. Liquid cultures of *C. difficile* were prepared by inoculating degassed BHIS broth with a single *C. difficile* colony and incubated overnight.

Aliquots (200 µl) of overnight *C. difficile* cultures were spread onto pre-reduced BHIS plates and incubated for 7 days to allow for sporulation. *C. difficile* strain 630 derivatives did not sporulate efficiently on BHIS and were therefore sporulated onto pre-reduced 70:30 agar plates instead (6.3% Bacto peptone, 1.1% Brain heart infusion, 0.4% protease peptone, 0.2% yeast extract, 0.03% L-cysteine, 9.8 mM Tris base, 5.3 mM (NH_4_)_2_SO_4_).

The resulting bacterial lawns were harvested by flooding the plates with ice-cold DIH_2_O. The samples were pelleted by centrifugation at 8,800 x g and the supernatants were decanted.

This wash procedure was repeated three times. Spores were then purified by density centrifugation through a 20% to 50% Histodenz gradient. After discarding the Histodenz solution, the spore pellet was washed three times with DIH2O, resuspended in a sodium thioglycolate solution (0.5 g/l), and stored at 4 °C until further use. To determine spore purity, samples were visualized by phase contrast microscopy or stained using the Schaeffer-Fulton technique [39]. Spores used in all experiments were more than 95% pure.

### Activation of *C. difficile* spores for germination

Purified *C. difficile* spores were washed three times with DIH2O, then heat shocked at 68 °C for 30 min to kill any vegetative cells that may have been present within the spore suspension. This process also activates spores to become more responsive to germinants [40]. After spores were heat shocked, samples were cooled to room temperature, and washed an additional three times with DIH2O. Spores were finally diluted to an optical density at 580 nm (OD_580_) of approximately 1.0 with sodium phosphate buffer (88 mM NaH_2_PO_4_, 12 mM Na_2_HPO_4_, pH 6.0), supplemented with 5 mg/ml sodium bicarbonate.

### Optical Density Assays

To test for *C. difficile* spore germination, 180 µl aliquots of the activated spore suspensions were added to 96-well clear plates containing taurocholate and glycine, at a final concentration of 6 mM and 12 mM, respectively. Following the addition of the spore suspension, the OD580 was measured every minute for 2 hours. As a negative control, 180 µl aliquots of spore suspension were treated with neat DMSO. All experimental conditions were performed in triplicate and the final volume in each well was 200 µl.

### Sequencing of *cspC* genes

Genomic regions flanking the *cspC* gene from *C. difficile* strains CDC38, DH1834, R20291, 05-1223-046, 630, 7004578, 8085054, and 9001966 were amplified to generate roughly 1.8 kbp amplicons. The PCR products were purified, and DNA was quantified by Nanodrop. All *cspC* amplicons were sequenced by Genewiz Sequencing, the DNA sequences were translated to protein sequences, and aligned through Benchling.

### Construction of *cspA, cspB* and *cspC* null mutants

*ΔcspA, ΔcspB*, and *ΔcspC* mutant *C. difficile* strains were constructed using allele-coupled exchange (ACE) protocols [20, 41, 42]. Briefly, upstream and downstream homology regions from each gene were PCR amplified and cloned into pMTL-YN3 by Gibson Assembly to generate pMTL-YN3:cspA, pMTL-YN3:cspB, and pMTL-YN3:cspC knockout constructs. These plasmids were transformed into *E. coli* strain Mach1 2T1 and their identity was confirmed by sequencing. Plasmids were then transformed into *E. coli* CA434.

*E. coli* CA434 harboring the pMTL-YN3:cspA, pMTL-YN3:cspB, or pMTL-YN3:cspC knockout constructs were individually cultured with shaking at 225 rpm at 37 °C for 6 h in Luria-Bertani (LB) medium, supplemented with chloramphenicol (25 µg/ml) and ampicillin (100 µg/ml). After centrifugation of 1 ml cultures, each *E. coli* pellet was transferred into the anaerobic workstation and gently resuspended in 1 ml of *C. difficile* strain 630*Δerm ΔpyrE* strain grown for 5 h in BHIS broth. The conjugation mating mixtures were then spotted onto pre-reduced BHIS plates lacking taurocholate. Following 16 h incubation, the conjugation mating mixtures were harvested, and the cell suspensions were spread onto pre-reduced BHISTKC plates (BHIS agar containing 10 µg/ml thiamphenicol, 50 µg/ml kanamycin, and 8 µg/ml cefoxitin).

The largest transconjugants that grew on BHISTKC were restreaked three consecutive times onto BHISTUCK15 (BHISTKC with thiamphenicol increased to 15 µg/ml). Transconjugants grown after three rounds of BHISTUCK15 were picked and restreaked onto *C. difficile* minimal medium (CDMM) plates supplemented with 5 µg/ml uracil and 2 mg/ml 5-fluoroorotic acid (CDMMUFOA). Individuals that grew on CDMMUFOA were restreaked and screened by PCR for the correct gene deletion and for wildtype contamination.

### Effect of Csp mutants on the hamster model of CDI

Female weaned golden Syrian hamsters were injected via IP with clindamycin as previously reported [34]. Hamsters (n=5 per group) were then challenged by oral gavage with 100 CFUs of wildtype *C. difficile* strain 630 spores or 1,000 CFUs of *C. difficile ΔcspA* mutant, *C. difficile ΔcspB* mutant, or *C. difficile ΔcspC* mutant spores. Animals were monitored for sign of disease twice daily. Any animal showing signs of distress was culled.

### Effect of CspC and bile salts on *C. difficile* strain 630 spore viability

Wildtype *C. difficile* strain 630 spores or *C. difficile ΔcspC* mutant spores were serially diluted and 100 µl from selected dilutions were plated onto BHIS, BHIS supplemented with 1.0 mM taurocholate, or BHIS supplemented with 10 µM CaPA. Plates were incubated anaerobically for 24-48 hours. Colony forming units (CFUs) were enumerated from plates showing non-confluent *C. difficile* colonies.

### Effect of bile salts on *C. difficile* strain R20291 spore viability

Activated *C. difficile* strain R20291 spores were serially diluted and 100 µl from selected dilutions were plated onto BHIS agar, BHIS agar supplemented with 1.0 mM taurocholate, BHIS agar supplemented with 10 µM CaPA, BHIS agar supplemented with 10 µM taurocholate, or BHIS agar supplemented with 100 µM CamSA. CFUs were determined as above.

### DPA release assay

Individual aliquots of activated *C. difficile* strain R20291 spores were supplemented with 12 mM glycine, and 35 µl of TbPV (250 µM TbCl3, 250 µM pyrocatechol violet, 10 mM HEPES, pH 7.0). Each aliquot was then supplemented with either DMSO, 1.0 mM taurocholate, 10 µM taurocholate, or 10 µM CaPA. At different time points, released DPA was measured by exciting the DPA-Tb complex at 270 nm and following the fluorescence emission increase at 545 nm on a Tecan Infinite M200 96-well plate reader (Tecan Group, Männedorf, Switzerland). An identical aliquot of *C. difficile* spores (OD ~ 1.0) was boiled for 2 hours to release all core Ca-DPA, thus representing maximum DPA release. Percent germination was determined by dividing each DPA signal by the DPA signal from boiled spores [43].

### Effect of CaPA concentration on *C. difficile* spore viability

Activated *C. difficile* strain R20291 spores were serially diluted and 100 µl from selected dilutions plated onto BHIS agar supplemented with 3.1, 6.2, 10, 20, or 40 µM CaPA. CFUs were determined as above.

### Enrichment of *C. difficile* individuals able to germinate with CaPA

Activated *C. difficile* strain R20291 spores were serially diluted and 100 µl from selected dilutions were plated onto BHIS agar supplemented with 1.0 mM taurocholate, or BHIS supplemented with 10 µM of CaPA. Colonies that grew on BHIS containing 10 µM of CaPA were pooled into 5 ml of degassed BHIS broth and plated for re-sporulation. The resulting spores were purified and re-plated as above. The enrichment process was repeated for four rounds. The percent of CFU recovery in CaPA plates was determined by dividing the CFUs from CaPA-plates by the CFUs from taurocholate-plates in each enrichment round.

### Effect of pre-germination on *C. difficile* spore viability

Activated *C. difficile* strain R20291 spore aliquots were pre-germinated by incubating for 30 minutes with either buffer, DMSO, 6 mM taurocholate + 12 mM glycine, or 6 mM taurocholate + 12 mM glycine + 10 µM CaPA. Spore samples were heat-treated at 68 °C for 30 minutes to eliminate germinated cells. Spores were serially diluted and 100 µl from selected dilutions were then plated onto BHIS agar supplemented with 1.0 mM taurocholate. CFUs were determined as above.

### Effect of direct taurocholate/CaPA competition on *C. difficile* spore viability

Activated *C. difficile* strain R20291 spores were serially diluted and 100 µl from selected dilutions were plated onto BHIS agar, BHIS agar supplemented with 1.0 mM taurocholate, BHIS agar supplemented with 1.0 mM taurocholate/10 µM CaPA, or BHIS agar supplemented with 1.0 mM taurocholate/50 µM CaPA. CFUs were determined as above.

### Effect of bile salt pre-incubation on *C. difficile* spore viability

Activated *C. difficile* strain R20291 spores were pre-bound with bile salts by incubating for 30 minutes in buffer containing increasing concentrations of either CaPA or taurocholate. As a control, separate spore aliquots were incubated with DMSO. Spores were serially diluted and 100 µl from selected dilutions were plated onto BHIS supplemented with increasing concentrations of taurocholate or CaPA. CFUs were determined as above.

## RESULTS

To investigate whether *cspC* variants could be correlated with differential spore responses to bile salts, we amplified and sequenced the *cspC* gene from eight different *C. difficile* strains. These strains were selected since they showed varied germination responses to taurocholate and chenodeoxycholate [22], as well as CamSA [33].

Sequence alignment showed that the CspC protein product is highly conserved, with only five SNPs differences between all strains tested (Fig. S1). Only two amino acid substitutions were found at each one of these SNPs. Four of the SNPs coded for amino acids that are structurally conserved and are predicted to have low mutation severity (http://www.insilicase.com/Web/SubstitutionScore.aspx).

The CspC protein sequence clustered into two groups. The CspC variant of strains 630 (ribotype 012), 8085054 (ribotype 014), and 9001966 (ribotype 106) showed 100% sequence identity. Similarly, three strains of ribotype 027 (R20291, CDC38, and DH1834) had 100% CspC sequence identity. The identity of the CspC variants were not correlated with the response of each strain to bile salts during the germination process [22].

When strain 630 was used as the reference strain, CspC from strains R20291, CDC38, and DH1834 showed four unique mutations (L38V, E151Q, V179A, and N394S). Strain 05-1223-046 exhibited the most divergent CspC but was more closely related to the R20291 CspC isoform. When aligned against strain R20291, CspC from strain 05-1223-046 exhibited a single G279W difference.

To determine the effect of CspABC complex on germination, we generated the corresponding *ΔcspA, ΔcspB*, and *ΔcspC* mutants on *C. difficile* strain 630 [20]. As previously reported, none of the *Δcsp* mutants were able to germinate in response to taurocholate and glycine when monitored by optical density assays (Fig. S2).

Since *C. difficile* spore germination is a prerequisite for disease, we also tested the ability of these *Δcsp* mutants to trigger CDI in antibiotic-treated hamsters. Whereas hamsters succumb to disease when administered approximately 100 wildtype *C. difficile* spores [34], hamsters receiving a 1,000 spores from *Δcsp* mutants did not exhibit any clinical signs of CDI (Fig. S3).

We further tested the effect of bile salts on spore viability of the *ΔcspC* mutant. As previously reported [44], the absence of bile salts leads to poor germination and low CFU recovery, even for wildtype spores. As expected, millimolar concentrations of taurocholate, the natural *C. difficile* spore germinant, allowed us to quantitatively recover and calculate for the total spore CFUs (Fig. 1). In fact, BHIS plates containing taurocholate increase CFU recovery of wildtype *C. difficile* strain 630 by three orders of magnitude over BHIS alone. Interestingly, supplementation of BHIS plates with CaPA resulted in a 16-fold CFU recovery compared to unsupplemented Bplates.

**Figure 1.**
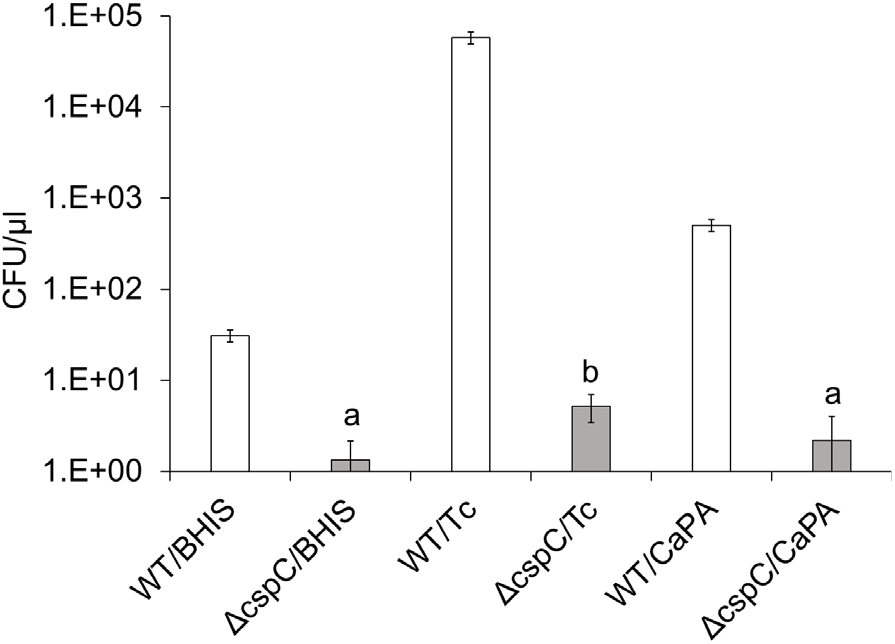
CspC mutant show compromised spore viability. (A) Wildtype *C. difficile* strain 630 spores (white columns) and *C. difficile* strain 630 *ΔcspC* spores (grey columns) were plated onto BHIS agar alone, BHIS agar supplemented with 1.0 mM taurocholate (Tc), or BHIS agar supplemented with 10 µM CaPA. Error bars represent standard deviations of at least three independent measurements. Columns that are labeled with different letters are statistically different (P > 0.05) by one-way ANOVA followed by Scheffé multiple comparison analysis.

On the other hand, the *ΔcspC* mutant exhibited severe germination deficiencies (Fig. 1, grey bars) compared to wildtype *C. difficile* strain 630 (Fig. 1, white bars), even in media that lacked bile salts. Supplementation of BHIS plates with taurocholate increased *ΔcspC* mutant CFU recovery four-fold, but still more than three order of magnitude lower than wildtype spore recovery. In contrast to wildtype spores, supplementation of BHIS plates with CaPA did not increase CFU recovery of *ΔcspC* mutant spores compared to unsupplemented BHIS plates.

To test the effect of bile salts on *C. difficile* strain R20291 CFU recovery and thus spore viability, equal amounts of spores were plated onto agar containing different bile salts. Similar to wildtype *C. difficile* strain 630 (Fig. 1), *C. difficile* strain R20291 shows four orders of magnitude increase in CFU recovery in the presence of taurocholate (Fig. 2A).

**Figure 2.**
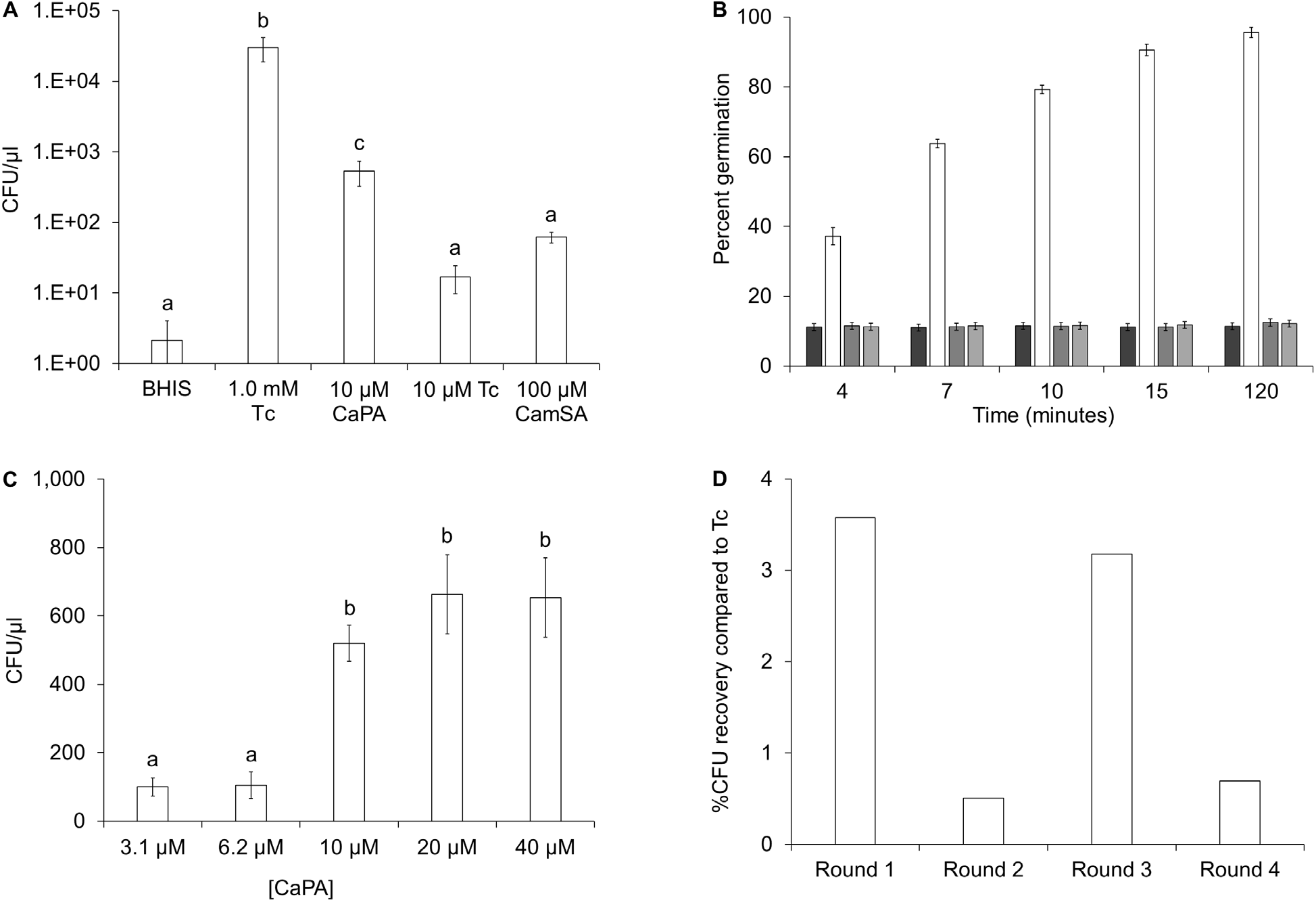
CaPA acts as a sub-optimal alternative germinant. (A) *C. difficile* strain R20291 spores were serially diluted, and aliquots were plated onto BHIS agar, BHIS agar supplemented with 1.0 mM taurocholate (Tc), BHIS agar supplemented with 10 µM CaPA, BHIS agar supplemented with 10 µM Tc, or BHIS agar supplemented with 100 µM CamSA. Plates were incubated anaerobically for 24-48 hours. Colony forming units (CFUs) were determined from plates showing non-confluent *C. difficile* colonies. (B) *C. difficile* strain R20291 spores were germinated in a solution containing 12 mM glycine and supplemented with either DMSO (dark grey bar), 6 mM taurocholate (white bar), 10 µM taurocholate (medium grey bar), or 10 µM CaPA (light grey bar). DPA release was monitored overtime by an increase in fluorescence emission of the Tb-DPA complex at 545 nm. Percent germination was determined by dividing each fluorescence measured by the fluorescence obtained from boiled spores. (C) *C. difficile* strain R20291 spores were plated into BHIS agar supplemented with 3.1 µM, 6.2 µM, 10 µM, 20 µM, or 40 µM CaPA. CFUs were determined as above. (D) *C. difficile* strain R20291 spores were plated onto BHIS agar supplemented with 10 µM of CaPA. The resulting colonies were pooled, re-sporulated and the resulting spores plated again onto BHIS agar supplemented with 10 µM of CaPA. The enrichment process was repeated for four rounds. The percent CFU recovery was determined by dividing the CFUs from CaPA-plates by the CFUs from taurocholate-plates in each enrichment round. In all panels, error bars represent standard deviations of at least three independent measurements. Columns that are labeled with different letters are statistically different (P > 0.05) by one-way ANOVA followed by Scheffé multiple comparison analysis.

Interestingly, micromolar concentrations of the CaPA anti-germinant leads to CFU recovery approximately two orders of magnitude above BHI alone (Fig. 2A). CFU recovery enhancement was not apparent when taurocholate was used at the same 10 μM concentration as CaPA. Furthermore CamSA, an anti-germinant with activity against *C. difficile* strain 630 but not *C. difficile* strain R20291, fails to increase CFU recovery above background, even when present at 10-fold higher concentrations than CaPA.

In contrast to the multi-day time frame of CFU recovery assays, DPA release tests the initial stages of spore germination. Spores treated with millimolar concentrations of taurocholate and glycine exhibited rapid DPA release that was completed within the first 15 minutes after germinant exposure (Fig. 2B). DPA release was not detected from spores incubated with glycine and treated with DMSO, 10 μM CaPA, or 10 μM taurocholate, even after two hours of incubation with germinant mixtures.

To determine the specificity for CaPA to affect CFU recovery, we serially increased CaPA concentrations in agar plates. Low concentrations of CaPA resulted in lower CFU recovery. CFU recovery was increased at higher CaPA concentrations, saturating at approximately 10 μM CaPA (Fig. 2C).

Since a subset of *C. difficile* strain R20291 spores were able to germinate in response to CaPA, we wanted to enrich for individuals able to recognize and utilize CaPA as a germinant, rather than as germination inhibitor. However, even after four sequential rounds of enrichment, CFU recovery with CaPA did not improve substantially and fluctuated between 0.5% and 3% compared to 1.0 mM taurocholate without showing an upward trend (Fig. 2D).

To determine if CaPA can prevent spore germination in the presence of taurocholate, spore suspensions were pre-treated with germinant mixtures, and heat shocked to kill newly germinated cells prior to plating onto taurocholate recovery plates. As expected, pre-treatment with DMSO had no effect on CFU recovery. Furthermore, pre-incubation with taurocholate and glycine resulted in 97% reduction of CFU recovery on taurocholate plates (Fig. 3). In contrast, addition of CaPA to the taurocholate/glycine germination mixture resulted in a 20-fold increase in CFUs on taurocholate-plates.

**Figure 3.**
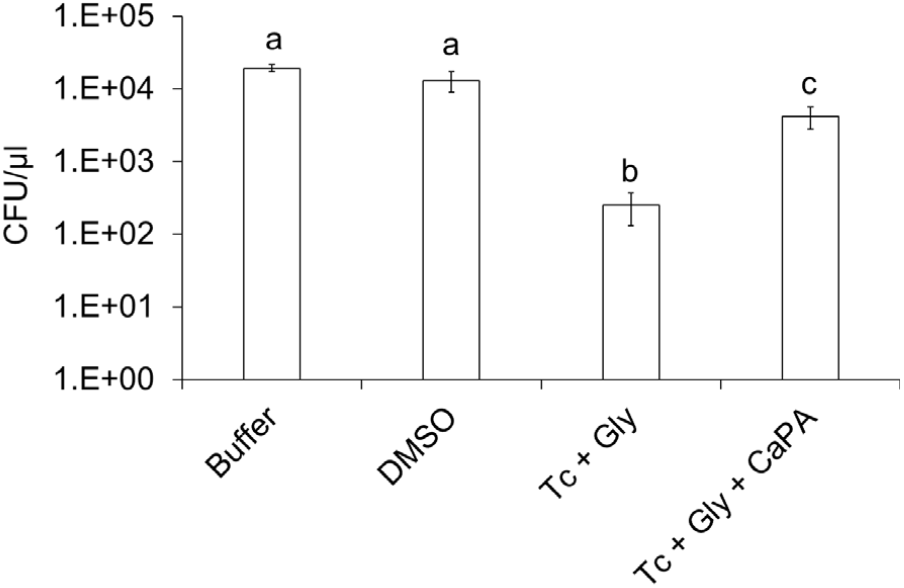
Effect of spore germination on CFU recovery. *C. difficile* strain R20291 spores were incubated with buffer, DMSO, taurocholate + glycine, or taurocholate + glycine + CaPA. Spore samples were then heat-treated to eliminate germinated cells. Spore aliquots were then plated onto BHIS agar supplemented with taurocholate. CFUs were determined as above. Error bars represent standard deviations of at least three independent measurements. Columns that are labeled with different letters are statistically different (P > 0.05) by one-way ANOVA followed by Scheffé multiple comparison analysis.

To test if CaPA could act as an inhibitor of taurocholate-mediated germination during a longer timeframe, we plated untreated *C. difficile* strain R20291 spores onto BHI agar containing both taurocholate and CaPA. Supplementation of taurocholate-plates with 10 µM CaPA reduced CFU recovery by approximately 30%, even though taurocholate is present at 100-fold higher concentration (Fig. 4A). Even more, when CaPA was increased to 50 µM, CFU recovery was reduced by approximately 85% compared to plates containing taurocholate alone.

**Figure 4.**
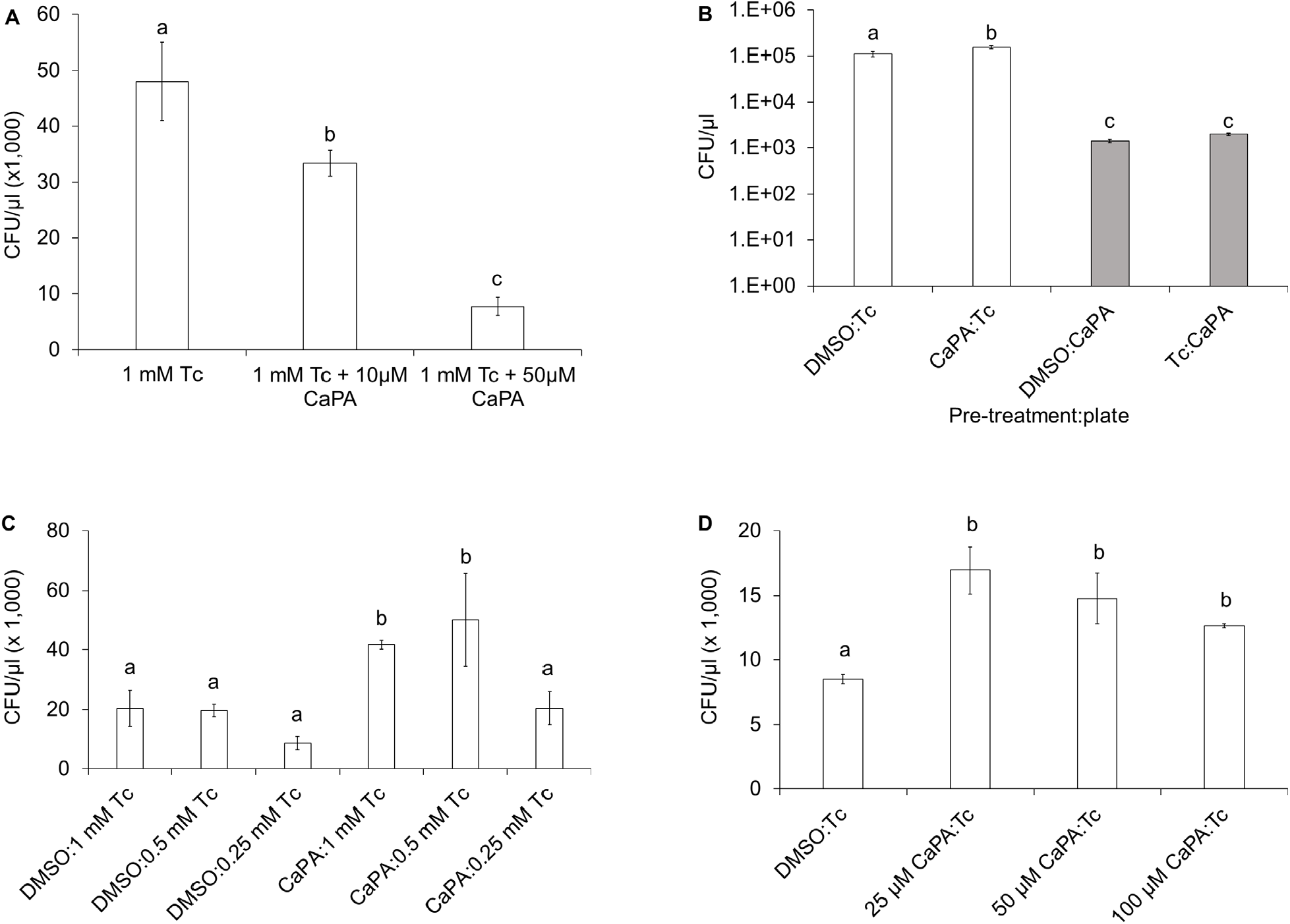
Effects of bile salt binding on CFU recovery. (A) *C. difficile* strain R20291 spores were plated into BHIS agar supplemented with 1.0 mM taurocholate (Tc), 1.0 mM Tc + 10 µM CaPA, or 1.0 mM Tc + 50 µM CaPA. (B) *C. difficile* strain R20291 spores were incubated in buffer containing either DMSO, Tc, or CaPA. Spore aliquots were then plated onto BHIS agar supplemented with either 1.0 mM Tc (white columns) or 10 µM CaPA (grey columns). (C) *C. difficile* strain R20291 spores were incubated in buffer containing either DMSO or 10 µM CaPA. Spore aliquots were then plated onto BHIS agar supplemented with either 1.0 mM Tc, 0.5 mM Tc or 0.25 mM Tc. (D) *C. difficile* strain R20291 spores were incubated in buffer containing either DMSO, 25 µM CaPA, 50 µM CaPA, or 100 µM CaPA. Spore aliquots were then plated onto BHIS agar supplemented with 1.0 mM Tc. In all panels, error bars represent standard deviations of at least three independent measurements. Columns that are labeled with different letters are statistically different (P > 0.05) by one-way ANOVA followed by Scheffé multiple comparison analysis.

To further test the antagonist/agonist effect of CaPA on *C. difficile* strain R20291 spore germination, spores were prebound with bile salts by incubating with either DMSO, taurocholate, or CaPA prior to plating into different bile salt containing plates. Pre-treatment with CaPA resulted in a statistically significant 40% increase in CFU recovery from taurocholate-supplemented plates compared to DMSO-treated controls (Fig. 4B, white columns). In contrast, pre-treatment with taurocholate did not affect CFU recovery in CaPA-supplemented plates compared to DMSO-controls (Fig. 4B, grey columns).

The increase of taurocholate-mediated CFU recovery by pre-incubation with CaPA is remarkable. We further tested this phenomenon by decreasing the concentration of taurocholate in the recovery plates. As before, pre-treatment with CaPA resulted in a doubling of CFUs at the higher concentrations of taurocholate. The CaPA enhancing effect was lost at the lowest taurocholate concentration tested (Fig. 4C). In contrast, increasing the concentration of the CaPA pre-treatment has no effect on CFU recovery on taurocholate-plates (Fig. 4D).

## DISCUSSION

We have targeted the germination process to prophylactically treat CDI. Our research has shown that bile salt analogs are able to inhibit *C. difficile* spore germination and, more importantly, prevent CDI in rodent models [33, 34, 36]. We recently found that structurally related bile salt analogs can inhibit spore germination in a strain-specific manner, even for strains that share taurocholate as their germination trigger [33]. To wit, CamSA (a cholate derivative with a *m*-sulfanilic acid side chain) was able to inhibit *C. difficile* strain 630 spore germination but failed against spores from the hypervirulent R20291 strain [36]. In contrast, CaPA (a cholate derivative with a simpler aniline side chain) was able to inhibit spore germination from multiple

*C. difficile* strains [33].

To understand these inter-strain differences, we first sequenced the *cspC* gene of eight different *C. difficile* strains, representing five different ribotypes. These strains were selected for their differential responses to activators and inhibitors of germination [22]. We found that the CspC protein was well conserved across these strains (Fig. S1). Furthermore, we found no correlation between any SNPs and germination profiles between the tested strains. Indeed, while the CspC protein from strain 630 is identical to the CspCs from strains 7004578, 8085054, and 9001966 only spores from strain 630 were inhibited by CamSA [22].

To experimentally probe for potential anti-germinant binding sites, we attempted to find naturally occurring *C. difficile* strain R20291 mutants resistant to CaPA inhibition using CFU recovery assays. Instead, we found that CaPA can trigger germination in up to 3% of *C. difficile* strain R20291 spores (Fig. 2A), a feat that was not obtained with CamSA, even when tested at 10-fold higher concentrations. This level of CFU recovery was unexpected since the frequency is well above the natural bacterial mutation rate (1:10^6^) and could suggest that a large subset of *C. difficile* spores is able to actively germinate in response to CaPA. However, we could not enrich the population of spores that respond to CaPA as a germinant.

Even though CaPA can mediate CFU recovery, it slows down germination and reduces spore viability. Consistent with previous data from optical density assays [33], CaPA treatment prevents rapid taurocholate/glycine-mediated germination onset (Fig. 2B and 3). This suggest that, similar to our previous analyses, CaPA binds tightly to *C. difficile* spores resulting in the inhibition of taurocholate-mediated spore germination [33]. Overall, CaPA-mediated germination was much more inefficient than taurocholate-mediated germination (Fig. 1 and 2A), showed concentration dependence (Fig. 2C), and seems to be stochastic instead of genetic-based (Fig. 2D). This is consistent with CaPA acting as an alternative substrate inhibitor of *C. difficile* spore germination [45].

The activity of CaPA as an alternate substrate inhibitor seems to depend on the order of binding to *C. difficile* spores. Simultaneous exposure to CaPA and taurocholate should allow spores to randomly bind these competing bile salts. Instead, long term incubation of spores in CaPA/taurocholate plates resulted in reduced CFU recovery and thus spore viability, even when the taurocholate concentration is orders of magnitudes higher than CaPA (Fig. 4A). This shows that CaPA can easily outcompete taurocholate for spore binding and is consistent with the strong inhibitory activity of CaPA [27].

In contrast, when taurocholate and CaPA are bound sequentially to *C. difficile* spores, we observed two different outcomes. Pre-treatment with CaPA prior to plating onto media containing taurocholate, surprisingly, resulted in an increase in CFU recovery (Fig. 4B-4D). This was unexpected since upon plating, CaPA would have been diluted over 100-fold and should have been released from any bile salt binding site. The fact that CaPA pre-treatment can affect spore viability for days after dilution suggests that CaPA irreversibly conditions spores for sub-optimal germination.

Even though pre-incubation with CaPA leads to increased CFU recovery in taurocholate plates, germination rates under these conditions remain compromised. Indeed, prior work have shown that CaPA pre-incubation results in strong germination inhibition in the shorter time frame of optical density assays [27, 33]. Thus, even though spore viability increases with CaPA-preincubation, their outgrowth rate is slow.

Interestingly, pre-treatment with taurocholate prior to dilution into CaPA-plates does not alter CFU recovery (Fig. 4B). This suggests that contrary to CaPA, taurocholate cannot lock spores in a pre-activated form to affect long-term spore viability.

These results are consistent with the existence of at least two distinct binding sites for bile salts in *C. difficile* spores [11]. This is further supported by the fact that a *ΔcspC* mutant can partially respond to taurocholate, but not to CaPA in CFU recovery assays (Fig. 1), suggesting that there is a secondary taurocholate-binding protein besides CspC *(i*.*e*. CspA). Although the nature of these distinct binding sites is not yet known, their presence would explain the inter-strain differences in anti-germination activity [18, 22] and CFU recovery (Fig. 2A) of CaPA and CamSA.

## CONCLUSIONS

Although Ger receptor homologs have been found in most sporulating bacteria, these receptors are conspicuously absent from *C. difficile*’s genomes [46]. However, given the physiological importance of the germination process, *C. difficile* must encode proteinaceous machinery that provides a link between the external environment and the initiation of germination.

In this study, we show that CspC variants do not correlate with anti-germinant selectivity by *C. difficile* strains. We further present evidence that CaPA is an alternative substrate inhibitor of *C. difficile* spores [47]. Even under conditions that allow increase spore viability, the presence of CaPA slows down the germination rate significantly. Hence, by binding tightly to *C. difficile* spores, CaPA might prevent optimal taurocholate-mediated spore activation. Instead, CaPA diverts spores into an alternative, suboptimal, and slowed down germination that reduces the number of spores able to enter the vegetative life cycle. Thus, toxin-producing *C. difficile* cells are never abundant enough to pose a risk of infection [33]. This process can account for the CDI prophylaxis activity of CaPA in animals.

Although not as well-known as other inhibition mechanisms, substrate inhibition is the most common deviation from classical kinetics, occurring in approximately 25% of known enzymes [48]. We have previously shown that both germination activation and inhibition can be approximated using Michaelis-Menten approaches in *Bacilli* and *Clostridia* spores [49-51].

Furthermore, classical kinetic analysis was used to infer putative germination receptors in *C. difficile* spores [12]. Hence, it is not surprising that CaPA inhibition of *C. difficile* spore germination follows a similar pathway.

CaPA is also a convenient molecular probe to further refine the mechanism of bile salt recognition by *C. difficile* spores. Our previous kinetic-based model assumed a homodimeric receptor with two identical binding sites that recognize taurocholate sequentially and cooperatively [11]. Based on published literature [8, 18-21] together with the data generated by our CFU recovery assays, we now hypothesize that CaPA binds tightly to spores in a CspC-dependent manner. Interaction with CaPA leaves spores irreversibly primed for sub-optimal germination. Taurocholate can then enter a second, separate bile salt binding site. Activation of the second bile salt binding site then allows *C. difficile* spores to recognize glycine. The net outcome of CaPA binding is a hobbled germination process that proceeds at a much slower pace and results in fewer *C. difficile* spores committed to returning to vegetative life.

It is possible that CaPA binds directly to CspC. However, we cannot rule out that CaPA could be binding to a separate unknown site upstream of CspC, thus inhibiting the signal cascade that triggers spore germination through the CspABC complex. Similarly, it seems that taurocholate can either bind to CspC, CspA, or even another unknown bile site receptor.

Given the importance of *C. difficile* spore germination and its relevance to the onset of CDI, there is a dire need to identify the receptors involved in bile acid recognition. By solving the identity of the bile acid receptor, a potential link could be established between germination phenotype and genes encoding the receptor(s).

## ACKNOWLEDGMENTS

This work was supported by the National Institute of Health [grant number R01-AI109139]. The authors thank Prof. Steve Firestine from Wayne State University for the synthesis of CaPA, Prof. Nigel Minton from University of Nottingham for providing clinical *C. difficile* strains, and Prof. Aimee Shen from Tufts University for the training and materials for *C. difficile* knock-out system.

## SUPPLEMENTARY FIGURES

**Figure S1.**
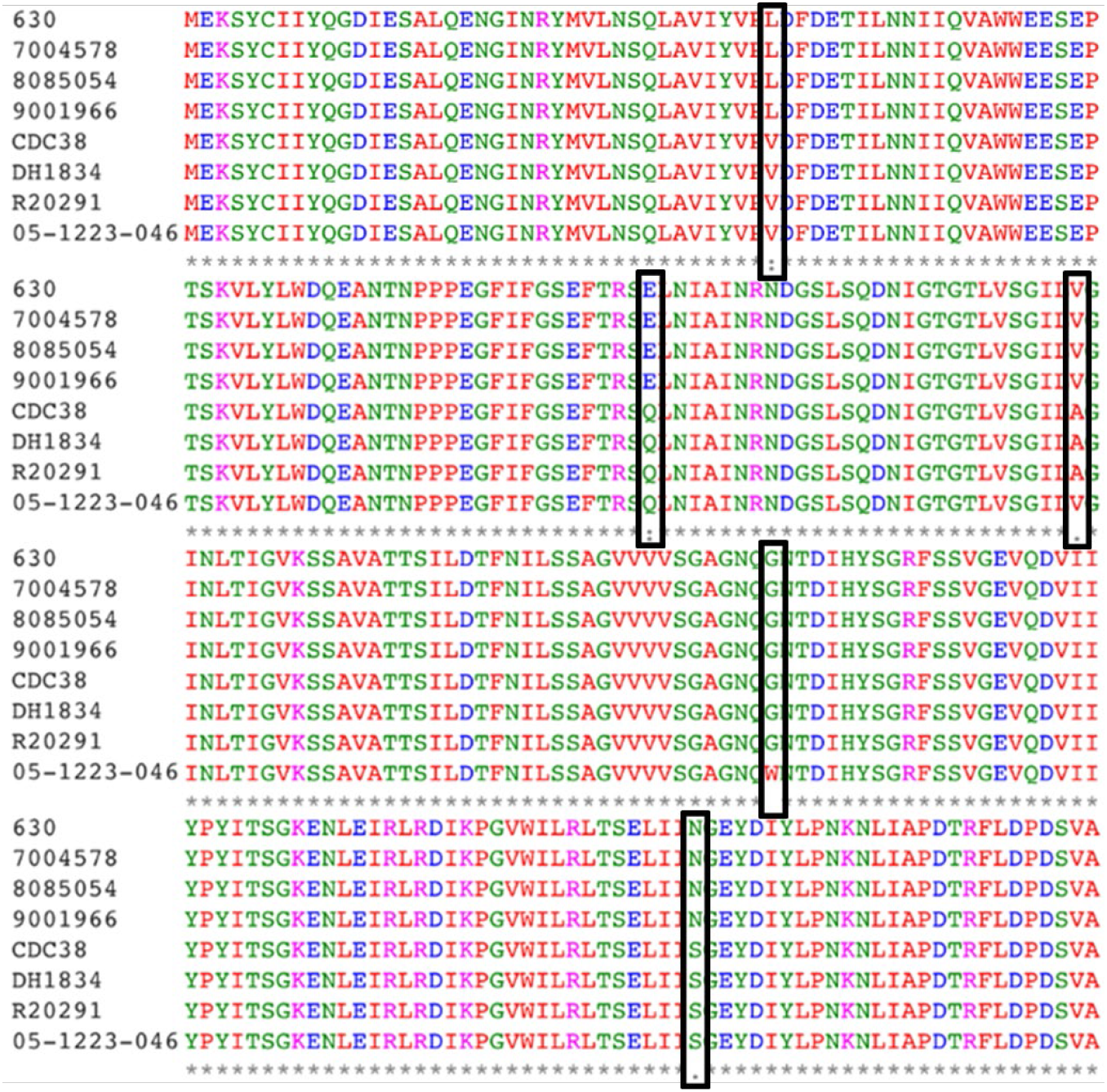
CspC sequence alignment from C. difficile strains with different germination bile salt profiles. The *cspC* gene was amplified from *C. difficile* strains 630, 7004578, 8085054, 9001966, CDC38, DH1834, R20291, and 05-1223-046. The nucleotide sequences of the *cspC* genes were then translated into protein sequences and aligned. Black boxes show positions with inter-strain amino acid variability.

**Figure S2.**
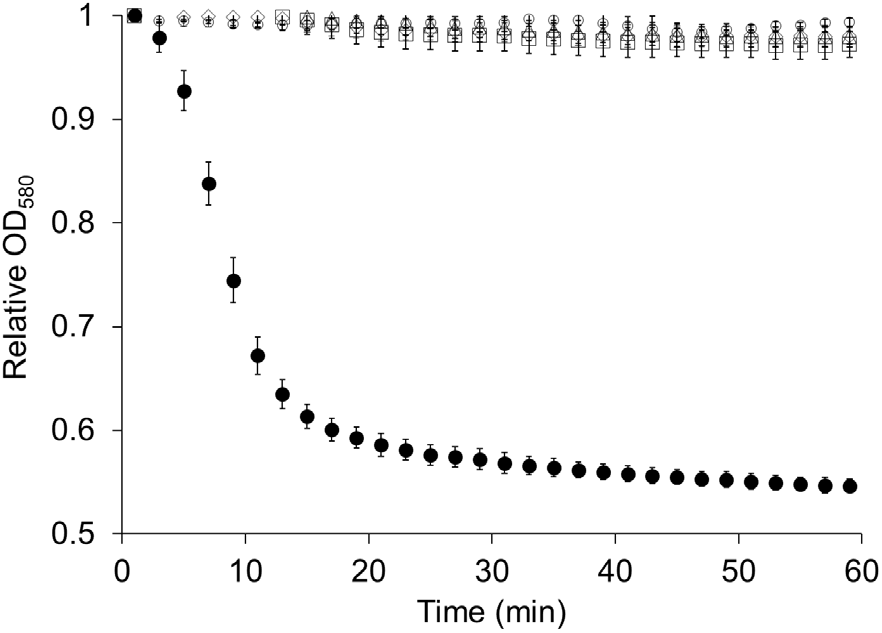
The cspABC operon is required for germination. *C. difficile* strain 630 wildtype spores (black circles), *ΔcspA* spores (white triangles), *ΔcspB* spores (white squares), and *ΔcspC* spores (white circles) were incubated with 6 mM taurocholate and 12 mM glycine. Germination was followed by the decrease of optical density at 580 nm (OD580). For clarity, data are shown at 5-min intervals. Error bars represent standard deviations of at least three independent measurements.

**Figure S3.**
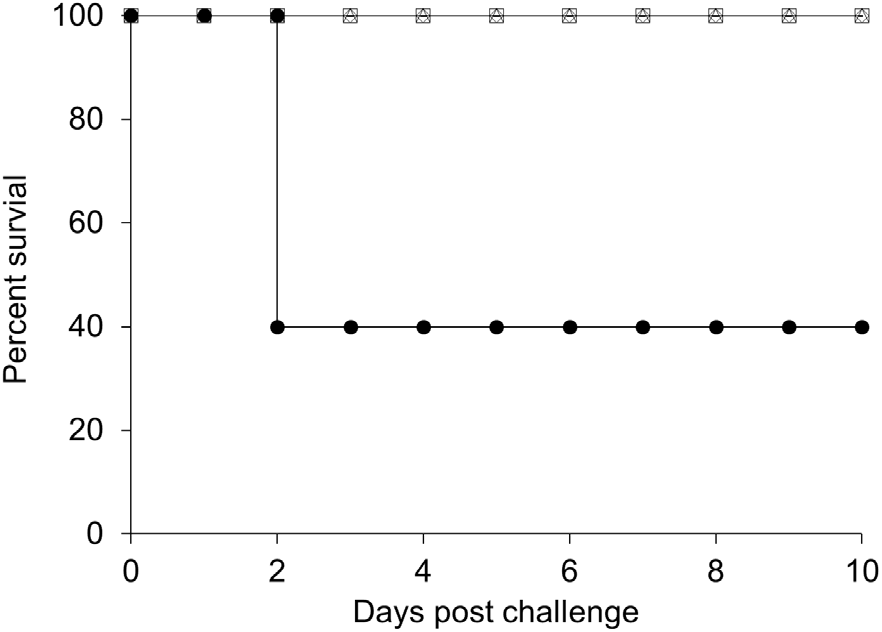
The cspABC operon is required for virulence in the hamster model of CDI. Kaplan-Meier survival plot of hamsters treated with 30 mg/kg clindamycin. Animals were challenged with 100 spores of *C. difficile* strain 630 wildtype (black circles), or/ 1,000 spores of either *ΔcspA* (white triangles), *ΔcspB* (white squares), or *ΔcspC* (white circles).

